# The failure of B cells to induce non-canonical *MYD88* splice variants correlates with lymphomagenesis via sustained NF-κB signaling

**DOI:** 10.1101/2020.06.18.154393

**Authors:** Yamel Cardona Gloria, Stephan H. Bernhart, Sven Fillinger, Olaf-Oliver Wolz, Sabine Dickhöfer, Jakob Admard, Stephan Ossowski, Sven Nahnsen, Reiner Siebert, Alexander N.R. Weber

## Abstract

Gain-of-function mutations of the TLR adaptor and oncoprotein MyD88 drive B cell lymphomagenesis via sustained NF-κB activation. In myeloid cells, sustained TLR activation and NF-κB activation lead to the induction of inhibitory *MYD88* splice variants that restrain prolonged NF-κB activation. We therefore sought to investigate whether such a negative feedback loop exists in B cells. Analyzing *MYD88* splice variants in normal B cells and different primary B cell malignancies, we observed that *MYD88* splice variants in transformed B cells are dominated by the canonical, strongly NF-κB-activating isoform of *MYD88* and contain at least three novel, so far uncharacterized signaling-competent splice isoforms. TLR stimulation in B cells unexpectedly reinforces splicing of NF-κB-promoting, canonical isoforms rather than the ‘MyD88s’, a negative regulatory isoform that is typically induced by TLRs in myeloid cells. This suggests that an essential negative feedback loop restricting TLR signaling in myeloid cells at the level of alternative splicing, is missing in B cells, rendering B cells vulnerable to sustained NF-κB activation and eventual lymphomagenesis. Our results uncover *MYD88* alternative splicing as an unappreciated promoter of B cell lymphomagenesis and provide a rationale why oncogenic *MYD88* mutations are exclusively found in B cells.

## Introduction

MyD88 has long been studied as an adaptor molecule for Toll-like receptor (TLR) and Interleukin-1 receptor (IL-1R) signaling in innate immunity ^1^. Its pivotal role is strikingly illustrated by the fact that loss-of-function mutations lead to severe immunodeficiency, whereas gain-of-function mutations promote oncogenesis : For example, rare dysfunctional alleles of *MYD88* compromise formation of the MyD88-mediated post-receptor complex ^2^, the so-called Myddosome ^3,4^, a pre-requisite for effective activation of the IL-1R-associated kinases (IRAKs) 2 and 4 and eventual activation of NF-κB and mitogen activated protein (MAP) kinases ^1^. Patients carrying loss-of-function *MYD88* alleles fail to respond to microbial TLR agonists and IL-1 and thus do not mount a sufficient innate immune response against pyogenic bacteria, leading to insufficient immunity and frequent premature death ^5^. Conversely, MYD88 mutations leading to constitutive Myddosome assembly ^6^, most notably the mutation Leu 265 to Pro (L265P) ^7^, are oncogenic and associated with sustained NF-κB signaling. L265P drives lymphoproliferation in murine models {Knittel, 2016 #334}. In humans, L265P is highly prevalent in various B cell malignancies {Ngo, 2011 #6882} but absent in other, e.g. myeloid, hematopoietic malignancies. Its strict occurrence in B cell malignancies has highlighted L265P’s diagnostic, chemo- and immunotherapeutic potential ^8–10^ but also posed the questions why only B cells are vulnerable to *MYD88* gain-of-function mutations? Additionally, the varying frequency of the L265P mutation in different B cell malignancies has been puzzling: Although the MyD88 L265P mutation may be found in up to 90% of Waldenström’s Macroglobulinemia patients ^11^, in diffuse large B cell lymphoma (DLBCL) and chronic lymphocytic leukemia (CLL) only 30 or 4% of patients carry this or other known gain-of-function *MYD88* mutations, depending on subtype ^7,12^. Thus, other mechanisms apart from mutation of *MYD88* appear to operate in L265P-negative patients, whereas a consistent “NF-κB signature” has been recognized as a unifying feature for most of these B cell malignancies ^13–15^.

The activation of NF-κB is also a primary outcome of MyD88-dependent signaling in myeloid cells ^1^. However, negative feedback on NF-κB signaling by alternative splicing operates in myeloid cells: TLR stimulation with LPS leads to the upregulation of a novel splice variant, then termed ‘MyD88 short’ (MyD88s, here also referred to as isoform 3, see Fig. 1A, B and Table 1) ^16^. Conversely to constitutive splicing ^17^, alternative splice variants arise from “alternative” splice sites in pre-mRNAs, that trigger, for example, exon skipping, alternative 5’ or 3’ splice sites within exon or intron sequences or intron retention. The resulting transcripts may be subject to frame shifts, premature termination codons and/or non-sense mediated decay (NMD) ^17,18^. Collectively, >90% of human multi-exon genes are subject to alternative splicing which expands the diversity and function of the proteome ^19,20^. In eukaryotes the spliceosome, where so-called splice factors (SFs) cooperate with five small nuclear ribonucleoprotein complexes (U1, U2, U4/U6, and U5), recognizes and assembles on introns to cleave and ligate RNA molecules for intron removal, generating protein-coding mRNAs ^21^. The spliceosome catalyzes splicing with high precision, but also displays high flexibility to regulatory signals for rapid responses, such as alternative splicing. Such a direct link between regulatory signals and innate immunity was recently proposed for the SF3A and SF3B mRNA splicing as both factors were shown to connect TLR signaling with the regulation of MyD88s ^22,23^.

**Table 1:** *MYD88* splice isoforms. Reference IDs from Ensembl and NCBI. ENST: cDNA sequence, ENSP: protein sequence, NM: curated NCBI mRNA, Protein-coding transcript; NP: NCBI protein coding sequence; Strep-HA: Strep III - Hemagglutinin tag; n/a: not available. *Values were calculated using ExPASy. ^§^ Generated constructs use Met1 as start codon.

| MyD88<br>isoforms | mRNA |  | Protein |  |  | Expression construct<br>MW (incl. Strep-HA<br>tag; kDa)* |
| --- | --- | --- | --- | --- | --- | --- |
|  | Reference ID | CDS<br>(bp) | Reference ID | Length (aa) | MW (kDa) |  |
| Isoform 1 | ENST00000421516.3<br>NM_001172567.2 | 915 | ENSP00000391753<br>NP_001166038.2 | 304 | 34.1 | 40.5 |
| Isoform 2 | ENST00000396334.8<br>NM_002468.5 | 891 | ENSP00000379625<br>NP_002459.3 | 296 | 33.0 | 37.4 |
| Isoform 3<br>(MyD88s) | ENST00000417037.7<br>NM_001172568.2 | 795 | ENSP00000401399<br>NP_001166039.2 | 264 | 29.6 | 34.7 |
| Isoform 4 | ENST00000651800.1<br>NM_001172569.3 | 615 | ENSP00000499012<br>NP_001166040.2<br>Uniprot: Q99836-3 | 204 | 22.1 | 27.2 |
| Isoform 5 | ENST00000650112.1<br>NM_001172566.2 | 480 | ENSP00000497991<br>Uniprot: Q99836-4 | 159 | 17.2 | 22.8 |
| Isoform 6 | ENST00000652213<br>NM_001365876.1 | 738 | ENSP00000498576<br>NP_001352805.1 | 245 | 27.1 | 34.9<sup>§</sup> |
| Isoform 7 | NM_001365877.1 | 642 | NP_001352806.1 | 213 | 23.5* | 29.9<sup>§</sup> |
| Isoform 8 | ENST00000652590.1 | 723 | n/a (new) | 240 | 26.4* | 32.8<sup>§</sup> |

**Figure 1:**
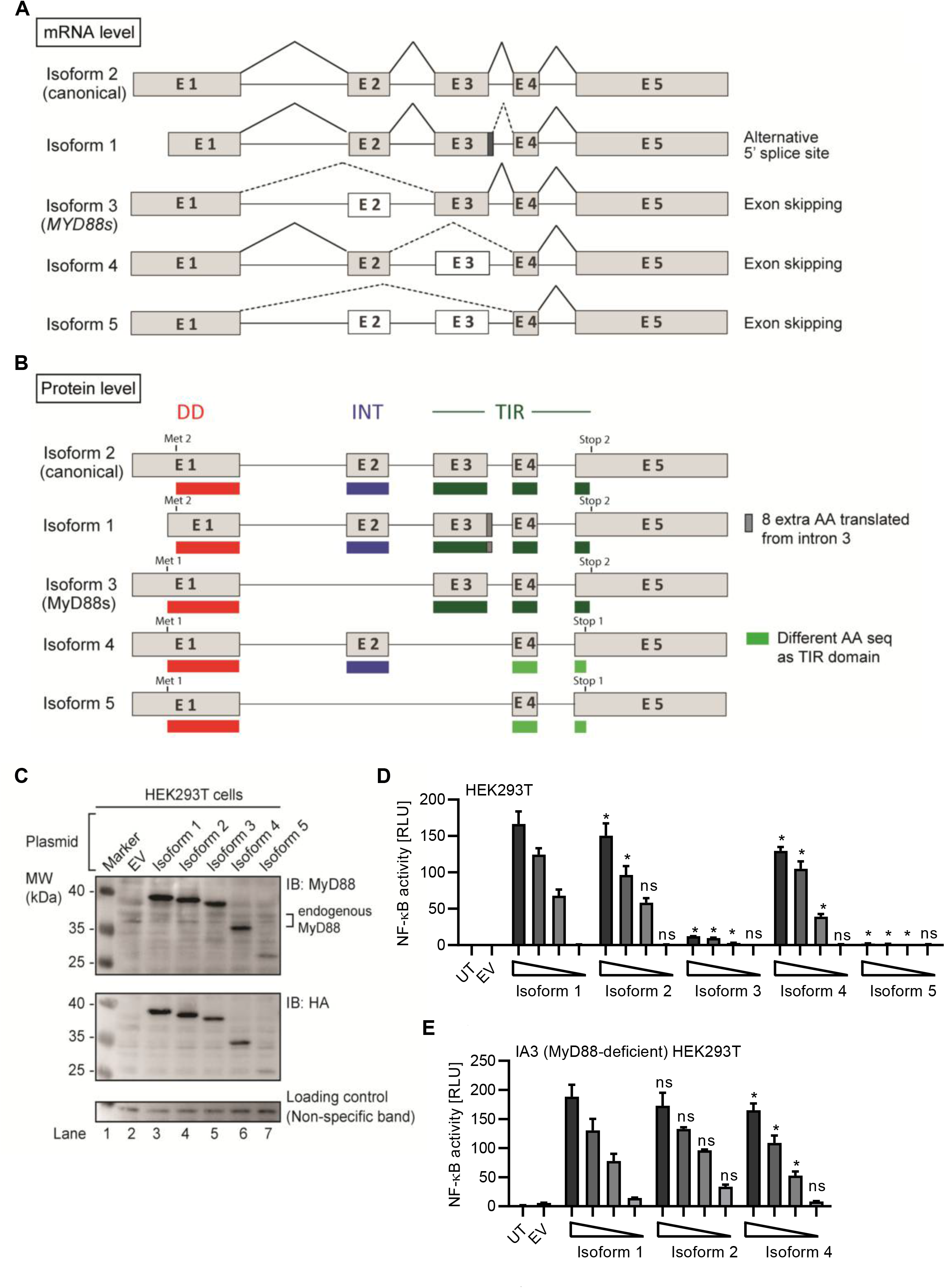
Several alternative MyD88 isoforms support NF-κB signaling. (A-B) Schematic representation of *MYD88* isoforms on mRNA (A) and protein (B) level. (C-E) HEK293T cells were transfected with plasmids for different *MYD88* splice isoforms and lysates analyzed for expression or pathway activation by immunoblot (C, n=3) or NF-κB dual luciferase assay (D, n=4), respectively. (E) as in D but using MyD88-deficient I3A cells (n=3). In C-E one representative of ‘n’ technical replicates is shown. * = p<0.05 according to two-way ANOVA comparing to isoform 1 (D, E).

MyD88s (isoform 3) represents an alternatively spliced in-frame deletion of exon 2 and thus a MyD88 variant significantly shorter than the canonical isoform 2: Whereas isoforms 1 and 2 contain the canonical N-terminal death domain (DD), central intermediate domain (ID) for IRAK recruitment and C-terminal Toll/IL-1R (TIR) domain for TLR binding, MyD88s (isoform 3) lacks the ID. The ID has been proposed to couple activated TLRs to the IRAK-containing Myddosome and thus transduce the incoming signal ^24^. Hence, MyD88s is signaling incompetent. Even though its characterization has been limited to myeloid cells, MyD88s (isoform 3) has been considered a primary negative regulator of this pathway and part of an essential negative feedback loop induced upon extended TLR signaling in myeloid cells ^25^. Isoform 1, the first reference sequence described, represents the longest transcript and translated protein for MyD88 by taking an alternative donor splice site 24 nt downstream of exon 3, adding 8 amino acids within the TIR domain. Apart from isoforms 1-3, two additional splice isoforms of *MYD88* have since been described, namely, isoforms 4 and 5 (Fig. 1A, B and Table 1), whose properties have been less studied. Additionally, whether alternative splicing and feedback regulation is operable in other non-myeloid immune cells has not been addressed.

We speculated that if a negative feedback loop existed in B cells, TLR activation should also induce MyD88s (isoform 3) and thereby limit ongoing signaling. However, we show here that in B cells this feedback loop does not seem to operate; rather TLR stimulation drives the canonical, i.e. NF-κB promoting, isoform and thus does not restrict extended NF-κB activation by diverting transcripts to less signaling competent isoforms like MyD88s (isoform 3) as in myeloid cells. In line with this, primary B cell malignancies showed significantly higher degrees of the canonical *MYD88* splice isoform. Our data highlight a critical difference between myeloid and B cell regulation of *MYD88* splicing that provides an explanation for the susceptibility of B cells to oncogenic *MYD88* mutation.

## Methods

### Study participants and sample acquisition

All patients and healthy blood donors included in this study provided their written informed consent before study participation. Approval for use of their biomaterials was obtained by the local ethics committee at the University Hospitals of Tübingen, in accordance with the Declaration of Helsinki, applicable laws and regulations. Further details in Supplemental information.

### Isolation and stimulation of primary human immune cells

Peripheral blood mononuclear cells (PBMCs) from healthy donors were isolated from whole blood or buffy coats (University Hospital Tübingen Transfusion Medicine) using Ficoll density gradient purification, primary B cells from PBMCs using B Cell Isolation Kit II (Miltenyi Biotec, >90% purity by anti-CD19 staining). PBMCs were rested and stimulated with 100 ng/ml LPS (from *E. coli* K12, Invivogen) or 0.1 μM CpG 2006 (TIB MOLBIOL), B cells with 2.5 μg/ml CpG 2006 and 5 μg/mL anti-human IgM (Fc5μ, Jackson Immuno Research). Carboxyfluorescein-succinimidyl ester (CFSE, Life Technologies) was used to track cell proliferation. Flow cytometry (BD FACSCanto II) was analyzed using FlowJo PC version 10. Further details in Supplemental Information.

### Plasmid constructs

N-terminally StrepIII-Hemagglutinin tagged *MYD88* isoform expression constructs were based on the reference sequences listed in Table 1 and generated by gene synthesis (Genewiz) or PCR cloning and verified by DNA sequencing. Further details in Supplemental Information.

### Cell cultures

All HEK293T and DLBCL cell lines were described and cultured as previously ^6^. THP-1 WT and MyD88-deficient cells were a kind gift from V. Hornung, Gene Center, Munich. Further details in Supplemental Information.

### Dual Luciferase Assay

Dual luciferase assays (DLA) were described previously ^6^. Briefly, *MYD88* isoforms (1-100 ng), NF-κB firefly luciferase reporter (100 ng) and Renilla luciferase control reporter (10 ng) were transfected into HEK293T cells. Further details in Supplemental Information.

### SDS-PAGE and immunoblot

Cell lysates (RIPA buffer with phosphatase and protease inhibitors) were separated on 10% or 4%–12% SDS-PAGE. Proteins blotted onto nitrocelullose membranes were probed with anti-HA H3663 (Sigma-Aldrich), MyD88 4D6 (Thermo Fisher), MyD88 D80F5 and 3699 (CST), HRP-conjugated secondary antibodies (1:8000) and visualized using CCD-based ECL detection. Further details in Supplemental Information.

### Quantitative PCR

Upon total RNA isolation (RNeasy Mini Kit, Qiagen) and reverse transcription, qPCR reactions (20 ng cDNA, 0.3 or 1 μM primers (Table S1), 1x FastStart Universal SYBR Green Master Rox, Sigma) were performed and normalized to GAPDH expression. Further details in Supplemental Information.

### Lymphoma, CLL and ovarian cancer dataset analysis

RNAseq libraries for Burkitt’s Lymphoma (BL, n=20), Follicular Lymphoma (FL, n=80), Diffuse Large B cell Lymphoma (DLBCL, n=71), FL-DLBCL (n=15), naïve B cells (n=5) and germinal center B cells (n=5) were from the European genome-phenome database Chronic Lymphocytic Leukemia (CLL) RNAseq data (n=289) from the ICGC-CLL Consortium (https://dcc.icgc.org/releases) ^26,27^. Ovarian cancer RNAseq libraries (n=85) were from the ICGC/OV-AU project (Australian Ovarian Cancer Study, https://dcc.icgc.org/projects/OV-AU) ^28,29^. Details regarding the analysis are given in Supplemental Information.

### Statistic analysis

Experimental data was analyzed using Excel 2010 (Microsoft) and/or GraphPad Prism 6, 7 or 8 or in *R*, flow cytometry data with FlowJo 10. Normal distribution in each group was always tested using the Shapiro-Wilk test first for the subsequent choice of a parametric (ANOVA, Student’s t-test) or non-parametric (e.g. Friedman, Mann-Whitney U or Wilcoxon) test. p-values (α=0.05) corrected for multiple testing were then calculated in Prism. Values <0.05 were generally considered as statistically significant and denoted by * or # throughout. Comparisons were made to unstimulated control, unless indicated otherwise, denoted by brackets.

## Results

### *MYD88* displays comprehensive splicing leading to functionally disparate isoforms

Given the importance that the MyD88s splice variant has been ascribed in murine myeloid cells ^16,22^, we sought to conduct a systematic characterization of all known human *MYD88* splice variants. Until recently, five *MYD88* mRNA transcripts with differential splicing have been reported (Table 1, Fig. 1A), giving rise to five protein isoforms with different domain structure (Fig. 1B). Compared to the canonical isoform 2, isoform 1 features an additional 8 amino acids in frame between exon 3 and 4, i.e. in the TIR domain, due to the use of an alternative splice site (dark grey box and/or dashed lines in Fig. 1B, S1B). Isoform 3 lacks the ID (exon 2) but includes both DD and TIR domain and corresponds to the aforementioned MyD88s variant. Isoform 4 and 5 both lack the TIR domain entirely, due to frame-shifts resulting from the skipping of exon 3 (Fig. S1A). In terms of canonical MyD88 domains, isoform 4 thus is limited to a DD-ID protein followed by 36 C-terminal amino acids that bear no apparent similarity to any known proteins (Fig. S1A). In isoform 5, exon 2 is additionally skipped, thus resulting in a DD-only variant. In order to investigate functional differences, these isoforms were cloned into StrepHA-tagged expression constructs and their expression verified in transfected HEK293T cells. Evidently, all constructs could be detected as proteins of 40, 37, 35, 27 and 23 kDa (Fig. 1C, Table 1), albeit with different expressions levels. The shortest isoform, termed isoform 5, was barely detectable, indicating it may be less stable. Next, we assessed the ability of all isoforms to drive NF-κB activation using dual luciferase assays upon transfection of equal amounts of expression plasmids in HEK293T cells. Whilst this assay cannot report on the ability to transduce incoming TLR signals, it is well established to assess MyD88 downstream signaling potential ^2,6,7,30–32^. Here, isoform 2, the canonical MyD88 form, was the most active, followed by isoform 1 (Fig. 1D). Isoform 4 was also able to induce NF-κB activity, at slightly lower levels. Isoform 3 and 5 were not able to induce NF-κB activity, consistent with a lack of ID, which is required to assemble into a Myddosome and recruit IRAK4 ^4,31^. Since HEK293T cells endogenously express MyD88 isoform 2 at high levels (*cf.* Fig. 1C), we also conducted the experiment in the MyD88-deficient HEK293T-derivative cell line, I3A ^30^. An almost identical picture emerged (Fig. 1E). Since murine and human MyD88s (isoform 3) was described as a dominant-negative regulator of canonical MyD88 due to lack of the ID ^31,33^, we also tested whether isoforms 3 and 5 could block TLR signaling, e.g. via TLR5, in the HEK293T system, but this was not the case (Fig. S1C, D). Collectively, non-canonical MyD88 isoforms with an intact DD and ID (isoforms 1 and 4) are capable of transmitting downstream NF-κB activity and their expression may thus support the function of the canonical MyD88 (isoform 2), whereas isoforms 3 and 5 are inactive.

### B cells express all five *MYD88* splice isoforms

All analyses on *MYD88* splicing have so far focused on myeloid cells but MyD88 can play an oncogenic role in B cells via NF-κB signaling ^10^. We therefore next characterized the expression of the five isoforms in several ABC and GCB DLBCL cell lines using isoform-specific primers to distinguish isoforms 1/2 from other isoforms (Fig. S2A, B, Methods and Table 2). This confirmed the expression of isoforms 3, 4 and 5 at mRNA level in these cell lines (Fig. 2A). Using lysates of these ABC and GCB cell lines and an antibody directed against the DD, multiple MyD88-specific bands were visible (Fig. 2B). Taking into account the predicted molecular weights of the alternative isoforms (Table 1) and their corresponding mRNA levels in BJAB cells vs primary B cells (*cf.* Fig. 2A), certain labeled bands in Fig. 2B are likely to correspond to isoform 3 and 5. This same pattern of bands was observed using a combination of 2 distinct anti-MyD88 antibodies (Fig. S2C). This suggests that B cells express multiple MyD88 splice isoforms both on mRNA and protein level.

**Figure 2:**
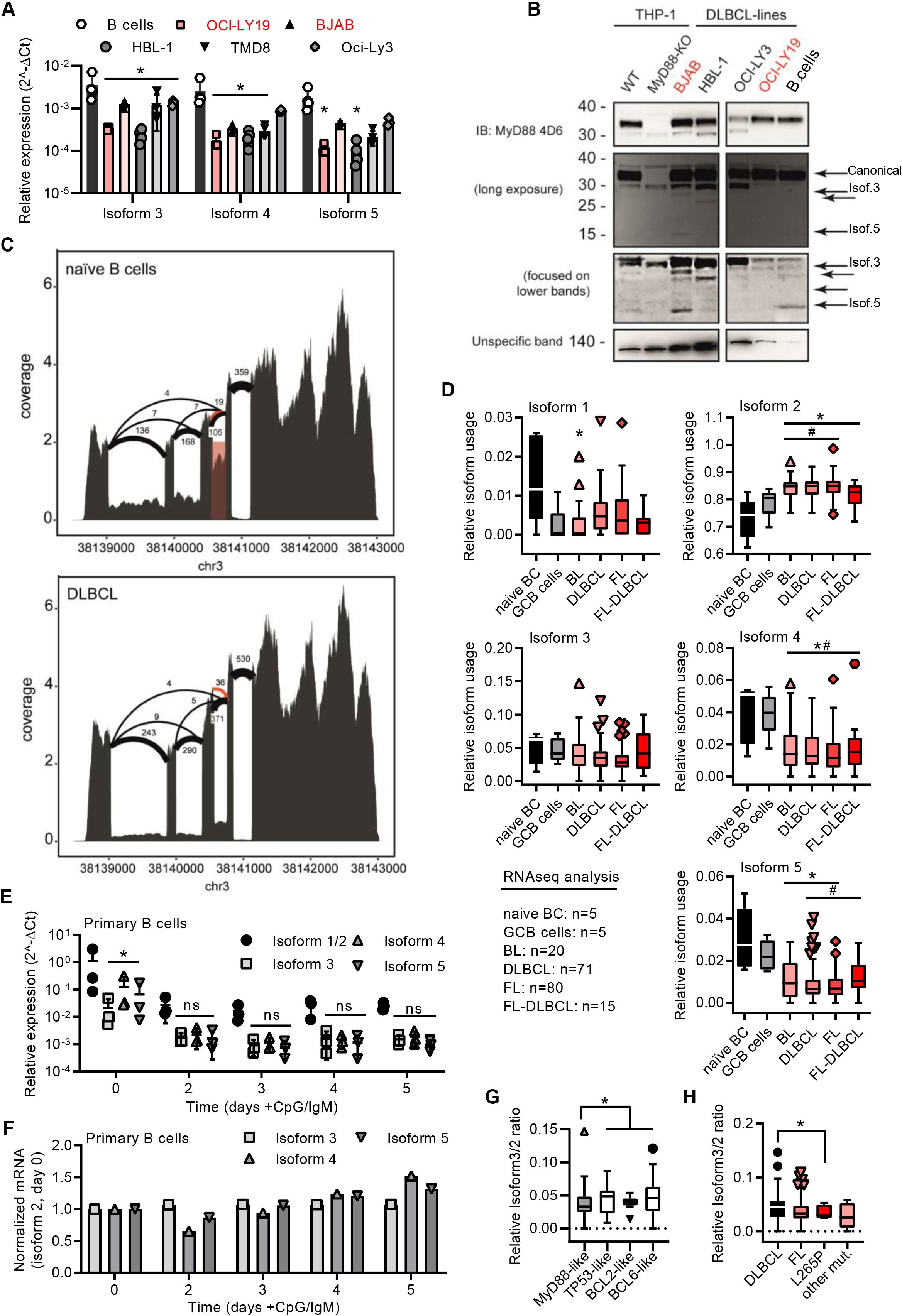
Lymphoma samples and stimulated B cells show a preference for the canonical *MYD88* isoform. (A) RT-qPCR analysis of isoforms 3-5 in primary B cells or lymphoma cell lines (n=3-4; red GCB, black ABC). (B) Immunoblot from THP-1 myeloid cells, B lymphoma cell lines and primary B cells (n=3). (C) Sashimi plots with mean read numbers supporting the splice junctions from naïve B cells (n=5) and DLBCL samples (n=83). Red shaded box shows intron retention and orange arcs represent an alternative donor splice site from isoforms 6 and 7. (D) RNAseq analysis of relative isoform usage from untransformed B cells or lymphoma samples (n=as indicated). Isoform 2 expressed as 1-(sum of all others). Other isoforms used: number of unique splice junctions divided by number of reads at the respective splice site (see Supplementary information). (E) RT-qPCR analysis of isoforms 1/2 - 5 in stimulated primary B cells (n=3). (F) as in E but normalized to isoform 2 at day 0. (G, H) Ratios of isoform 3 (*MYD88s*) to isoform 2 in different DLBCL sub-clusters (G) and in dependence of *MYD88* mutations (H). A and D-H represent combined data (mean+SD, or Tukey box and whiskers) from ‘n’ biological replicates (each dot represents one replicate). In B one representative of ‘n’ technical replicates is shown. * or # = p<0.05 according to Wilcoxon (G, H), Mann-Whitney (D, comparison to naïve B cells (*) or to GCB cells (#)), or two-way ANOVA (A, E and F).

As these transformed cells may not reflect primary tumors, we next characterized the expression of the five isoforms in primary B cell lymphoma samples and untransformed naïve B cells. Sashimi plots of RNAseq data from a total of 186 different lymphoma cases (Burkitt lymphoma, DLBCL, follicular lymphoma, follicular lymphoma-DLBCL), untransformed germinal center B cells (GCB, n=5) and naïve peripheral blood B cells (n=5, acquired by the German ICGC MMMLSeq consortium, see Methods) showed expression of all five isoforms at mRNA level (Fig. 2C, S2D). Interestingly, the canonical isoform 2 was significantly more abundant in transformed vs untransformed B cells, whereas other isoforms were either comparable between these groups (isoform 3) or significantly lower (isoform 1, isoform 4 and isoform 5) (Fig. 2D). Thus, transformed samples showed a preference for the canonical isoform 2 – but not isoform 3 (MyD88s) or other non-canonical isoforms. This was surprising as an ‘NF-κB signature’ has been attributed to these types of entities ^13–15^ and in myeloid cells NF-κB signaling was proposed to induce MyD88s (isoform 3). Collectively, this suggests that, contrary to expectations, lymphoma samples show a higher ratio of canonical MyD88 to MyD88s than naive B cells. Based on what has been published regarding the induction of MyD88s via NF-κB signaling in myeloid cells ^16,33^, we next tested whether defined NF-κB activating stimuli, e.g. TLR and BCR stimulation, would lead to an upregulation of isoform 3 in freshly purified (Fig. S2E) primary B cells. However, TLR9 CpG + IgM stimulation reduced *MYD88* transcription altogether and did not lead to higher relative induction of the MyD88s (isoform 3, Fig. 2E, F). In fact *MYD88* expression was generally downregulated, despite the fact that TLR stimulation was effective at driving cellular proliferation as assessed by CFSE proliferation assays (Fig. S2F). Conversely, when PBMC (which include monocytes, i.e. myeloid cells) were stimulated with LPS or CpG, MyD88s (isoform 3) was significantly upregulated (Fig. S2G), in line with earlier studies ^16,33,34^. Additionally, the analysis of sub-clusters (dependent on driver mutations) of DLBCL samples suggested that those driven by direct activators of NF-κB signaling (e.g. an *‘*MyD88-like’ sub-cluster, see Methods) had a lower ratio of alternative splicing vs canonical, and specifically isoform 3, than those driven by indirect NF-κB activation (e.g. BCL2-, BCL6- and TP53-like DLBCL, see Figs. 2G and S2H). In line with this, samples with NF-κB-promoting *MYD88* gain-of-function mutations, such as L265P, had a lower isoform 3 vs isoform 2 ratio, i.e. expressed significantly more isoform 2 vs isoform 3 transcripts (Fig. 2H). We conclude that proliferating B cells, like lymphoma samples, show and maintain a preference for canonical MyD88 signaling. Furthermore, in B cells NF-κB signaling does not induce or coincide with a shift towards inhibitory isoforms as reported for myeloid cells regarding MyD88s (isoform 3). Rather, the canonical, signaling-competent isoform 2 dominates.

### Novel MyD88 isoforms with TIR truncation in B cells are supportive of NF-κB signaling

In the process of RNAseq analysis we noticed additional alternative splicing events, namely either usage of another donor splice site within the exon 3 (leading to isoforms 6 and 7) or the retention of the exon 3-4 intron (here termed isoform 8), see Fig. 2C, 3A, B and Table 1. The novel splice site within exon 3 (20 nt upstream of a canonical donor) showed a Human Splicing Finder (HSF) score of 81. Typically, a score above 65 is considered a strong splice site ^35^, indicating these additional splicing events are highly plausible. This alternative donor site leads to a premature STOP codon and thus results in additional isoforms with a truncated TIR domain (Fig. 3A, B), which have not been reported so far. When expression constructs corresponding to isoforms 6-8 were transfected into HEK293T cells, proteins of the expected size (29 kDa for isoform 6, 24 kDa isoform 7 and 26 kDa for isoform 8; plus 6 kDa from the StrepHA-tag) were detectable (Fig. 3C and Table 1). To gain an insight into their ability to signal to NF-κB, we performed NF-κB dual luciferase assays in normal HEK293T and I3A cells as before. Evidently, isoforms 6 and 8 were able to induce downstream NF-κB activation in HEK293T cells, whereas isoform 7 did not (Fig. 3D, E). Isoform 6-8 transcripts were also detectable in the lymphoma samples (Fig. 3F-H) and, as with the other non-canonical isoforms, they were significantly less abundant in lymphoma cells vs naive B cells. In the 289 RNA-seq samples of the ICGC Chronic Lymphocytic Leukemia (CLL) dataset, 7 isoforms could be readily detected and quantified, with the canonical isoform showing the highest relative abundance, followed by isoform 6, while isoform 5 showed the lowest abundance (Fig. 3I). Furthermore, there were noticeable reads mapping to the exon 3-4 intron (Fig. S3C) confirming isoform 8 in CLL. All eight *MYD88* splice isoforms were also detectable in non-immune cells, as verified in a publicly available RNAseq dataset ^28^ for ovarian cancer (Fig. S3D). On the whole, there are 3 additional splice isoforms of MyD88 with truncated TIR domains out of which two unexpectedly can support signaling upon overexpression, similar to the canonical MyD88 isoform. This extended analysis highlights an even higher diversity of splice variants emanating from the *MYD88* oncogene than previously thought. Furthermore, splicing in B cell lymphomas appears to strongly favor the canonical *MYD88* isoform without diverting splicing events to alternative or signaling-incompetent splice isoforms. Importantly, we find no evidence for a significant induction of MyD88s (isoform 3) as a restrictor of TLR pathway activity.

**Figure 3:**
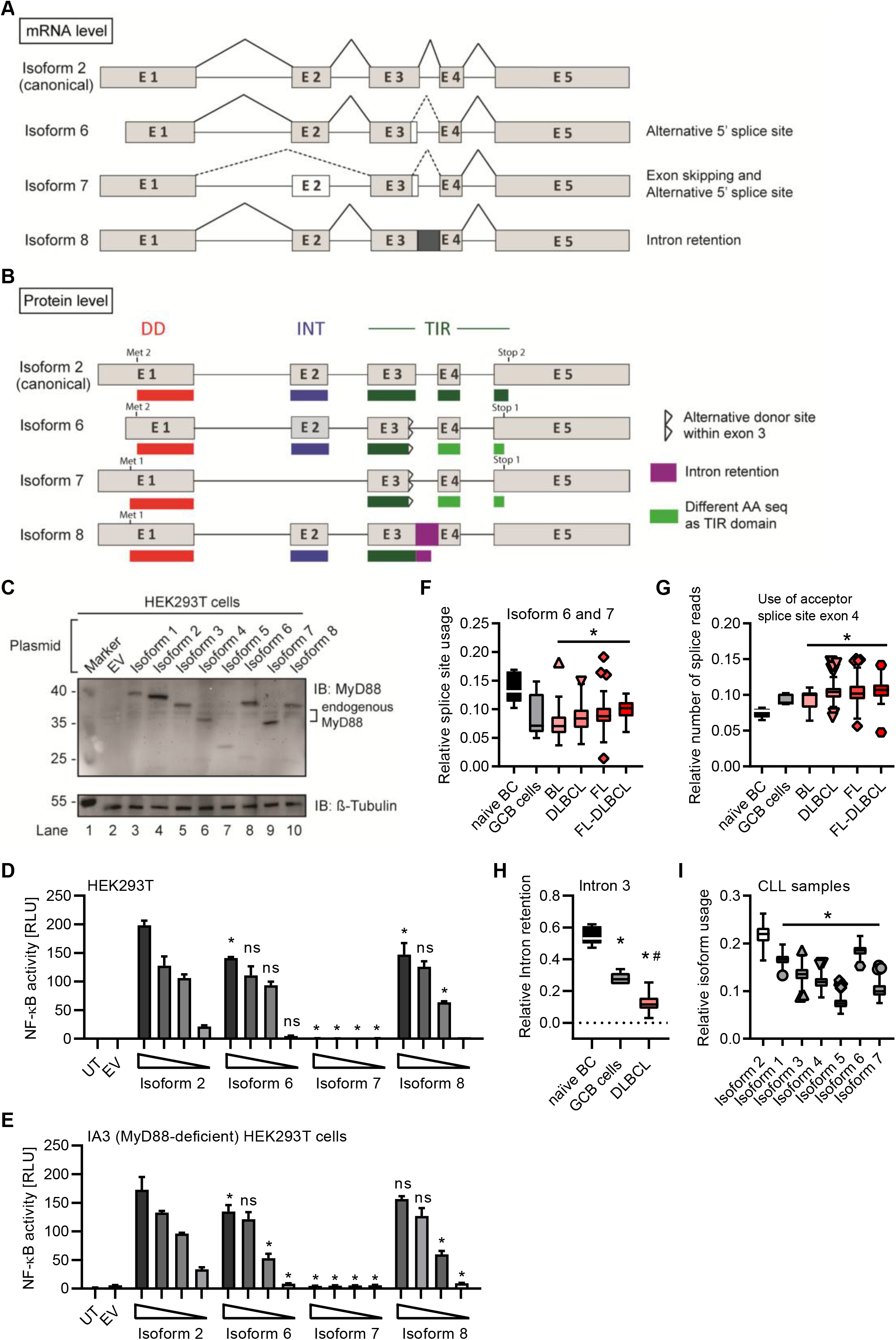
*MYD88* gives rise to three additional MyD88 isoforms. (A, B) Schematic representation of novel *MYD88* isoforms on mRNA and protein level. (C-E) HEK293T cells were transfected with plasmids for different *MYD88* isoforms and lysates analyzed for expression or pathway activation by immunoblot (C, n=2) or NF-κB dual luciferase assay (D, E n=3), respectively. (E) as in D but using MyD88-deficient I3A cells (n=3). (F-H) RNAseq analysis from untransformed B cells or lymphoma samples (n=as indicated). Intron retention presented as relative coverage of intron 3 compared to mean of flanking exons 3 and 4 (H). (I) RNAseq analysis from CLL samples (n=289). In C-E one representative of ‘n’ technical replicates is shown. F-I represent combined data (Tukey box and whiskers) from ‘n’ biological replicates (each dot represents one replicate). * or # = p<0.05 according to two-way ANOVA comparing to isoform 2 (D, E) or Wilcoxon Mann-Whitney (F-I) in comparison to naïve B cells (*, F-H) and to GCB cells (#, F-H) or isoform 2 (I).

## Discussion

Alternative splicing has emerged as a frequent phenomenon employed for fine-tuning or regulating signaling pathways and plays a pivotal role in the adaptive immune system ^36,37^. However, decisive regulators of innate immune pathways have also been subject to alternative splicing: Since its discovery in 2002, the induction of MyD88s via NF-κB signaling loop has been viewed as a classical example of an inflammation-restricting negative feedback loop in innate immunity ^16,25^. Hence, all the numerous subsequent studies on MyD88 splicing have exclusively focused on this isoform ^22,23,38–40^ and have been largely limited to myeloid cells, primarily in the murine system.

We here provide a comprehensive characterization of all currently reported human *MYD88* splice isoforms. This includes the novel isoforms 6-8, which are the only variants to contain partial TIR domains. During the course of this analysis, isoforms 6 and 7 were added to Genebank but had not been confirmed or studied in detail. Isoform 8 is a novel and surprisingly frequent splicing event not reported before and found abundantly in naïve B cells. Our analysis suggests that, with the exception of isoforms 3 (MyD88s), 5 and 7, isoforms (4, 6 and 8) may induce downstream NF-κB activity in overexpression assays. Whether they can nucleate or engage in the Myddosome in response to TLR signaling in the absence of a complete TIR domain remains to be studied. Potentially, isoforms 4, 6 and 8 may also be signaling incompetent. Thus all isoforms, except, isoforms 1 and 2, lead to dysfunctional MyD88 proteins. This would make our observations made on transcript levels even more striking as then none of the alternative splicing events would be able to counteract constitutive NF-κB signaling via isoform 2. Consequently, the oncogenic influence of isoform 2 is likely to be even more dominant.

Furthermore, we show that *MYD88* splicing is much more multi-faceted than previously reported: Our data indicate that whereas normal B cells use a richer repertoire of splice isoforms, the transformed status rather displays a reduced diversity and appears to suppress alternative splice events. The reason for this is unknown but our data warrant a further investigation in additional cohorts and entities, e.g. Waldemström’s macroglobulinemia, in future. Based on our data it appears that the preference for canonical isoform 2 and thus unrestricted NF-κB signaling may be favored in the oncogenic process. BCL2, BCL6 or TP53-driven lymphomas, which have an indirect effect on the NF-κB signature, showed lower levels of canonical *MYD88* and higher levels of isoform 1 and isoform 4, compared to MyD88-like lymphomas (Figs. 2G and S2H). This fits well with the observation that the gain-of-function mutation, L265P, leads to extended NF-κB hyperactivation and is a hallmark of oncogenic B cells ^7,41^. Of note, our data indicate that B cells lack the myeloid-specific negative feedback mechanism of MyD88s induction to rescue mutated cells from MyD88-driven oncogenesis: For example, TLR stimulation induced MyD88s in TLR-stimulated PBMC but not in B cells, and MyD88s was also not prominently expressed or regulated by typical NF-κB stimuli in B cell lymphoma cell lines and samples. Thus, B cells with increased NF-κB activity, due to L265P mutation or other mechanisms, cannot get “reigned in” (controlled) via MyD88s expression, unlike myeloid cells, and thus may support continued NF-κB pro-survival activity (Fig. 4). Our data thus provide an explanation why oncogenic mutations have only been reported in B cell lymphoma, rather than tumors arising from myeloid cells, whose MyD88s induction loop probably renders them more resistant to MyD88 pathway induced NF-κB activity.

**Figure 4:**
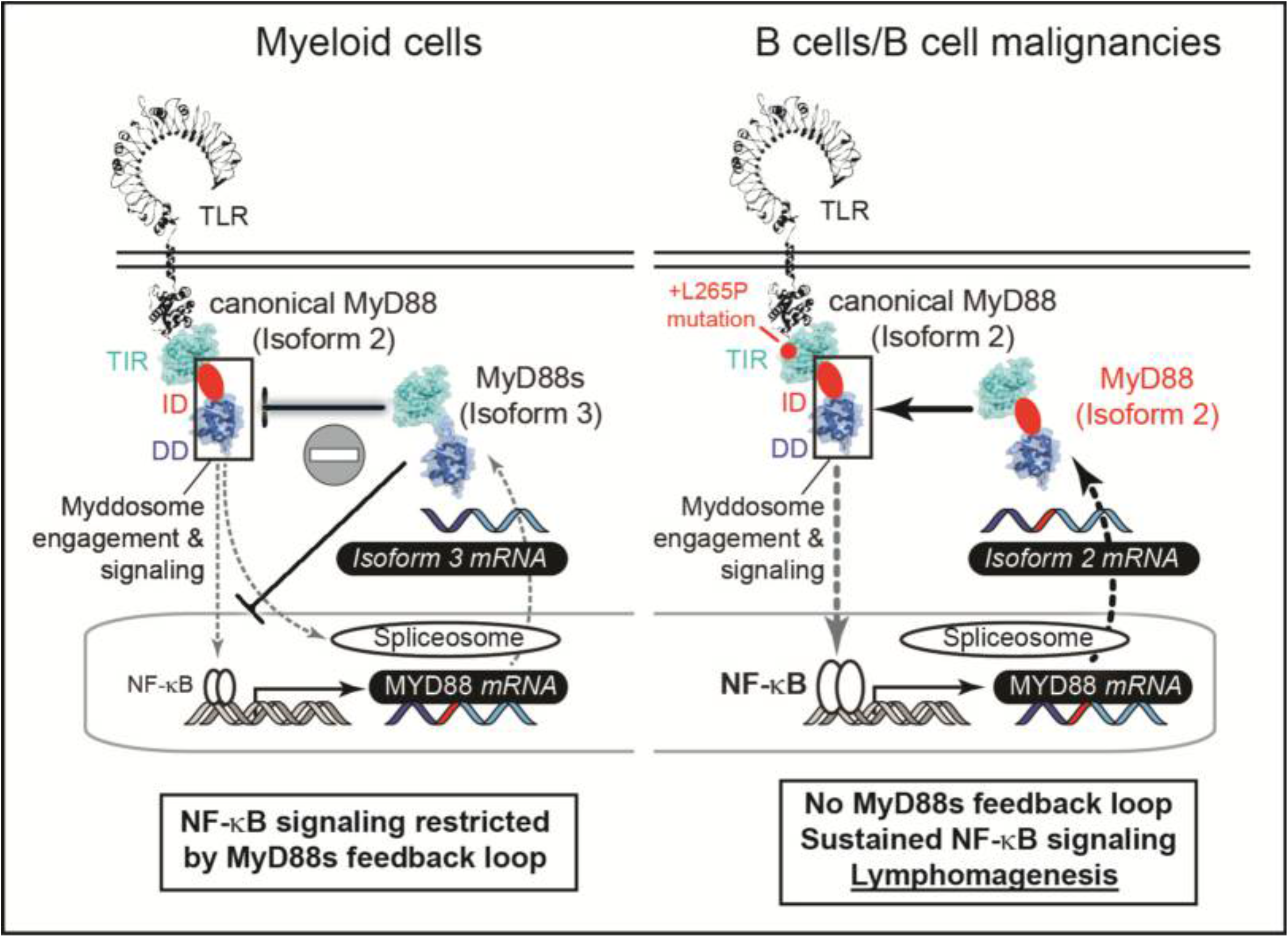
The failure of B cells to induce non-canonical *MYD88* splice variants correlates with lymphomagenesis via sustained NF-κB signaling. Graphical abstract summarizing splicing patterns in myeloid cells (left, previous work) and B cells (right, focus in this study).

Our observations that alternative splicing of genes in the MyD88 dependent pathway are important candidates in oncogenesis agrees with the recent description of oncogenic *IRAK4* isoforms, albeit in myeloid malignancies ^42^. It is intriguing to speculate whether the aforementioned negative feedback loop, that is absent in B cells, prevents MyD88 mutations from manifesting themselves but does not prevent oncogenic signaling arising from the next downstream pathway member, IRAK4. Undoubtedly, with the availability of powerful sequencing techniques the analysis of alternative splice isoforms of MyD88 pathway members for discovering novel non-mutational cancer drivers is both possible and warranted. In the substantial percentage of cases without druggable driver mutations this may offer opportunities for targeting e.g., via antisense oligonucleotide-mediated exon skipping ^43,44^. In this therapeutic sense, MyD88s or the other signaling incompetent isoforms described here may provide a blueprint for such an approach in B cell lymphomas.

## Acknowledgements

YCG, OOW and ANRW were supported by the Deutsche Forschungsgemeinschaft (DFG) SFB 685 ‘Immunotherapy’. YCG, OOW, SD, JA, SO, SF, SN and ANRW were also supported by the University of Tübingen Medical Faculty and the University of Tübingen. Infrastructural funding was provided by the University of Tübingen, the University Hospital Tübingen and the DFG Cluster of Excellence "iFIT – Image-Guided and Functionally Instructed Tumor Therapies" (EXC 2180, to AW). ANRW is a Deutsches Konsortium für Translationale Krebsforschung (DKTK; German Cancer Consortium) and Cluster of Excellence iFIT (EXC 2180) "Image-Guided and Functionally Instructed Tumor Therapies" PI. The authors gratefully acknowledge the Gauss Centre for Supercomputing e.V. (www.gauss-centre.eu) for funding this project by providing computing time on the GCS Supercomputer SuperMUC at Leibniz Supercomputing Centre (www.lrz.de).

## Author contributions

YCG, OOW and SD performed experiments; YCG, SHB, SF, SN, SD, JA, SO analyzed data; RS and SO were involved in sample collection; YCG and ANRW conceived and ANRW supervised the entire study; YCG and ANRW wrote the manuscript and all authors provided additions and comments to the manuscript.

## Conflict of interest

The authors declare that they have no conflict of interest.

## Abbreviations

BL: Burkitt lymphoma
CLL: chronic lymphocytic leukemia
DD: death domain
DLA: dual luciferase assay
DLBCL: diffuse large B cell lymphoma
FL: follicular lymphoma
GCB: germinal center B cells
HEK: human embryonic kidney
ID: intermediate domain
IL-1R: interleukin-1 receptor
IRAK: IL-1R-associated kinase
LPS: lipopolysaccharide
MyD88: myeloid differentiation 88
NF-κB: nuclear factor κB
TIR: Toll/Interleukin-1 receptor
TLR: Toll-like receptor
TNF: tumor necrosis factor
WT: wild-type

## Notes

### Competing Interest Statement

The authors have declared no competing interest.

## References

1. Kawai T, Akira S. Toll-like receptors and their crosstalk with other innate receptors in infection and immunity. Immunity. 2011;34(5):637–650.

2. George J, Motshwene PG, Wang H, et al. Two Human MYD88 Variants, S34Y and R98C, Interfere with MyD88-IRAK4-Myddosome Assembly. J Biol Chem. 2011;286(2):1341–1353.

3. Motshwene PG, Moncrieffe MC, Grossmann JG, et al. An oligomeric signalling platform formed by the toll-like receptor signal transducers MyD88 and IRAK4. J Biol Chem. 2009.

4. Lin SC, Lo YC, Wu H. Helical assembly in the MyD88-IRAK4-IRAK2 complex in TLR/IL-1R signalling. Nature. 2010;465(7300):885–890.

5. von Bernuth H, Picard C, Jin Z, et al. Pyogenic bacterial infections in humans with MyD88 deficiency. Science. 2008;321(5889):691–696.

6. Avbelj M, Wolz OO, Fekonja O, et al. Activation of lymphoma-associated MyD88 mutations via allostery-induced TIR-domain oligomerization. Blood. 2014;124(26):3896–3904.

7. Ngo VN, Young RM, Schmitz R, et al. Oncogenically active MYD88 mutations in human lymphoma. Nature. 2011;470(7332):115–119.

8. Nelde A, Walz JS, Kowalewski DJ, et al. HLA class I-restricted MYD88 L265P-derived peptides as specific targets for lymphoma immunotherapy. Oncoimmunology. 2017;6(3):e1219825.

9. Wilson WH, Young RM, Schmitz R, et al. Targeting B cell receptor signaling with ibrutinib in diffuse large B cell lymphoma. Nat Med. 2015;21(8):922–926.

10. Weber ANR, Cardona Gloria Y, Cinar O, Reinhardt HC, Pezzutto A, Wolz OO. Oncogenic MYD88 mutations in lymphoma: novel insights and therapeutic possibilities. Cancer Immunol Immunother. 2018;67(11):1797–1807.

11. Treon SP, Xu L, Yang G, et al. MYD88 L265P somatic mutation in Waldenstrom's macroglobulinemia. The New England journal of medicine. 2012;367(9):826–833.

12. Puente XS, Pinyol M, Quesada V, et al. Whole-genome sequencing identifies recurrent mutations in chronic lymphocytic leukaemia. Nature. 2011;475(7354):101–105.

13. Lenz G, Wright G, Dave SS, et al. Stromal gene signatures in large-B-cell lymphomas. The New England journal of medicine. 2008;359(22):2313–2323.

14. Thome M. CARMA1, BCL-10 and MALT1 in lymphocyte development and activation. Nat Rev Immunol. 2004;4(5):348–359.

15. Staudt LM. Oncogenic activation of NF-kappaB. Cold Spring Harb Perspect Biol. 2010;2(6):a000109.

16. Janssens S, Burns K, Tschopp J, Beyaert R. Regulation of interleukin-1- and lipopolysaccharide-induced NF-kappaB activation by alternative splicing of MyD88. Curr Biol. 2002;12(6):467–471.

17. Wang Y, Liu J, Huang BO, et al. Mechanism of alternative splicing and its regulation. Biomed Rep. 2015;3(2):152–158.

18. Ge Y, Porse BT. The functional consequences of intron retention: alternative splicing coupled to NMD as a regulator of gene expression. Bioessays. 2014;36(3):236–243.

19. Llorian M, Gooding C, Bellora N, et al. The alternative splicing program of differentiated smooth muscle cells involves concerted non-productive splicing of post-transcriptional regulators. Nucleic Acids Res. 2016;44(18):8933–8950.

20. Pimentel H, Parra M, Gee S, et al. A dynamic alternative splicing program regulates gene expression during terminal erythropoiesis. Nucleic Acids Res. 2014;42(6):4031–4042.

21. Wahl MC, Will CL, Luhrmann R. The spliceosome: design principles of a dynamic RNP machine. Cell. 2009;136(4):701–718.

22. De Arras L, Alper S. Limiting of the innate immune response by SF3A-dependent control of MyD88 alternative mRNA splicing. PLoS Genet. 2013;9(10):e1003855.

23. O'Connor BP, Danhorn T, De Arras L, et al. Regulation of toll-like receptor signaling by the SF3a mRNA splicing complex. PLoS Genet. 2015;11(2):e1004932.

24. Burns K, Janssens S, Brissoni B, Olivos N, Beyaert R, Tschopp J. Inhibition of interleukin 1 receptor/Toll-like receptor signaling through the alternatively spliced, short form of MyD88 is due to its failure to recruit IRAK-4. J Exp Med. 2003;197(2):263–268.

25. Liew FY, Xu D, Brint EK, O'Neill LA. Negative regulation of toll-like receptor-mediated immune responses. Nat Rev Immunol. 2005;5(6):446–458.

26. Ferreira PG, Jares P, Rico D, et al. Transcriptome characterization by RNA sequencing identifies a major molecular and clinical subdivision in chronic lymphocytic leukemia. Genome Res. 2014;24(2):212–226.

27. Puente XS, Bea S, Valdes-Mas R, et al. Non-coding recurrent mutations in chronic lymphocytic leukaemia. Nature. 2015;526(7574):519–524.

28. Patch AM, Christie EL, Etemadmoghadam D, et al. Whole-genome characterization of chemoresistant ovarian cancer. Nature. 2015;521(7553):489–494.

29. O'Donnell T, Christie EL, Ahuja A, et al. Chemotherapy weakly contributes to predicted neoantigen expression in ovarian cancer. BMC Cancer. 2018;18(1):87.

30. Li X, Commane M, Burns C, Vithalani K, Cao Z, Stark GR. Mutant cells that do not respond to interleukin-1 (IL-1) reveal a novel role for IL-1 receptor-associated kinase. Mol Cell Biol. 1999;19(7):4643–4652.

31. Janssens S, Burns K, Vercammen E, Tschopp J, Beyaert R. MyD88S, a splice variant of MyD88, differentially modulates NF-kappaB- and AP-1-dependent gene expression. FEBS Lett. 2003;548(1-3):103–107.

32. Loiarro M, Sette C, Gallo G, et al. Peptide-mediated interference of TIR domain dimerization in MyD88 inhibits interleukin-1-dependent activation of NF-{kappa}B. The Journal of biological chemistry. 2005;280(16):15809–15814.

33. Adib-Conquy M, Adrie C, Fitting C, Gattolliat O, Beyaert R, Cavaillon JM. Up-regulation of MyD88s and SIGIRR, molecules inhibiting Toll-like receptor signaling, in monocytes from septic patients. Crit Care Med. 2006;34(9):2377–2385.

34. Andrews CS, Miyata M, Susuki-Miyata S, Lee BC, Komatsu K, Li JD. Nontypeable Haemophilus influenzae-Induced MyD88 Short Expression Is Regulated by Positive IKKbeta and CREB Pathways and Negative ERK1/2 Pathway. PLoS One. 2015;10(12):e0144840.

35. Desmet FO, Hamroun D, Lalande M, Collod-Beroud G, Claustres M, Beroud C. Human Splicing Finder: an online bioinformatics tool to predict splicing signals. Nucleic Acids Res. 2009;37(9):e67.

36. Mourich DV, Iversen PL. Splicing in the immune system: potential targets for therapeutic intervention by antisense-mediated alternative splicing. Current opinion in molecular therapeutics. 2009;11(2):124–132.

37. Martinez NM, Lynch KW. Control of alternative splicing in immune responses: many regulators, many predictions, much still to learn. Immunol Rev. 2013;253(1):216–236.

38. Vickers TA, Zhang H, Graham MJ, Lemonidis KM, Zhao C, Dean NM. Modification of MyD88 mRNA splicing and inhibition of IL-1beta signaling in cell culture and in mice with a 2'-O-methoxyethyl-modified oligonucleotide. J Immunol. 2006;176(6):3652–3661.

39. Blumhagen RZ, Hedin BR, Malcolm KC, et al. Alternative pre-mRNA splicing of Toll-like receptor signaling components in peripheral blood mononuclear cells from patients with ARDS. Am J Physiol Lung Cell Mol Physiol. 2017;313(5):L930–L939.

40. Feng Z, Li Q, Meng R, Yi B, Xu Q. METTL3 regulates alternative splicing of MyD88 upon the lipopolysaccharide-induced inflammatory response in human dental pulp cells. J Cell Mol Med. 2018;22(5):2558–2568.

41. Knittel G, Liedgens P, Korovkina D, et al. B-cell-specific conditional expression of Myd88p.L252P leads to the development of diffuse large B-cell lymphoma in mice. Blood. 2016;127(22):2732–2741.

42. Smith MA, Choudhary GS, Pellagatti A, et al. U2AF1 mutations induce oncogenic IRAK4 isoforms and activate innate immune pathways in myeloid malignancies. Nat Cell Biol. 2019;21(5):640–650.

43. Cirak S, Arechavala-Gomeza V, Guglieri M, et al. Exon skipping and dystrophin restoration in patients with Duchenne muscular dystrophy after systemic phosphorodiamidate morpholino oligomer treatment: an open-label, phase 2, dose-escalation study. Lancet. 2011;378(9791):595–605.

44. Gramlich M, Pane LS, Zhou Q, et al. Antisense-mediated exon skipping: a therapeutic strategy for titin-based dilated cardiomyopathy. EMBO molecular medicine. 2015;7(5):562–576.

45. Lopez C, Kleinheinz K, Aukema SM, et al. Genomic and transcriptomic changes complement each other in the pathogenesis of sporadic Burkitt lymphoma. Nature communications. 2019;10(1):1459.

46. Hezaveh K, Kloetgen A, Bernhart SH, et al. Alterations of microRNA and microRNA-regulated messenger RNA expression in germinal center B-cell lymphomas determined by integrative sequencing analysis. Haematologica. 2016;101(11):1380–1389.

47. Kretzmer H, Bernhart SH, Wang W, et al. DNA methylome analysis in Burkitt and follicular lymphomas identifies differentially methylated regions linked to somatic mutation and transcriptional control. Nat Genet. 2015;47(11):1316–1325.

48. Richter J, Schlesner M, Hoffmann S, et al. Recurrent mutation of the ID3 gene in Burkitt lymphoma identified by integrated genome, exome and transcriptome sequencing. Nat Genet. 2012;44(12):1316–1320.

49. Hoffmann S, Otto C, Doose G, et al. A multi-split mapping algorithm for circular RNA, splicing, trans-splicing and fusion detection. Genome Biol. 2014;15(2):R34.

50. Katz Y, Wang ET, Silterra J, et al. Quantitative visualization of alternative exon expression from RNA-seq data. Bioinformatics. 2015;31(14):2400–2402.

51. Doose G, Bernhart SH, Wagener R, Hoffmann S. DIEGO: detection of differential alternative splicing using Aitchison's geometry. Bioinformatics. 2018;34(6):1066–1068.

52. Broseus L, Ritchie W. Challenges in detecting and quantifying intron retention from next generation sequencing data. Comput Struct Biotechnol J. 2020;18:501–508.

53. Sturm M, Schroeder C, Bauer P. SeqPurge: highly-sensitive adapter trimming for paired-end NGS data. BMC Bioinformatics. 2016;17:208.

54. Dobin A, Davis CA, Schlesinger F, et al. STAR: ultrafast universal RNA-seq aligner. Bioinformatics. 2013;29(1):15–21.

55. Katz Y, Wang ET, Airoldi EM, Burge CB. Analysis and design of RNA sequencing experiments for identifying isoform regulation. Nat Methods. 2010;7(12):1009–1015.

