## Supplementary information for "The failure of B cells to induce non-canonical *MYD88* splice variants correlates with lymphomagenesis via sustained NF-κB signaling"

**Title**

### Supplemental Methods

*Study participants and sample acquisition*

All patients and healthy blood donors included in this study provided their written informed consent before study participation. Approval for use of their biomaterials was obtained by the local ethics committee at the University Hospitals of Tübingen, in accordance with the principles laid down in the Declaration of Helsinki as well as applicable laws and regulations. Patient recruitment, sample acquisition and preparation of B cell lymphoma, CLL and ovarian cancer patients are described below. Healthy blood donors were recruited at the Interfaculty Institute of Cell Biology, Department of Immunology, University of Tübingen.

*Isolation and stimulation of primary human immune cells*

Peripheral blood mononuclear cells (PBMCs) from healthy donors were isolated from whole blood or buffy coats (University Hospital Tübingen Transfusion Medicine) using Ficoll density gradient purification. PBMCs were rested at least 4 h prior to stimulation, and resuspended and maintained for experiments in supplemented RPMI (Life Technologies). 5x10^6 PBMCs were stimulated with 100 ng/ml LPS (from *E. coli* K12, Invivogen) or 0.1 µM CpG 2006 (TIB MOLBIOL) for 0, 6, 12 and 18 h, then cells were lysed in RLT buffer + β-mercaptoethanol (Qiagen) for RT-qPCR measurements. Primary B cells were isolated from PBMCs by negative selection using B Cell Isolation Kit II (Miltenyi Biotec) according to instructions. B cell isolation always reached >90% purity and cells were seeded in supplemented RPMI with 10% human serum and rested for at least 4 h before the experiment. 5 x 10^6 B cells were stimulated for 0, 2, 3, 4, 5 days with 2.5 µg/ml CpG 2006 and 5 µg/mL anti-human IgM (Fc5µ, Jackson Immuno Research) and processed for RT-qPCR, or cells were additionally stained on day 0 with carboxyfluorescein-succinimidyl ester (CFSE, Life Technologies) to track cell proliferation. On day 5, proliferative cells were washed and stained with anti-CD19 Pacific Blue (Biolegend), then cells were analyzed on a BD FACSCanto™ II system with 488 nm excitation for CFSE and 405 nm for Pacific Blue. Graphs were generated using software FlowJo PC version 10.

*Plasmid constructs*

*MYD88* isoform expression constructs were generated in pTO-N-SH vector using the Gateway cloning system (Thermo Fisher) adding a fused StrepIII-Hemagglutinin tag at the N-terminus of the gene of interest. The coding sequences (CDS) of MyD88 isoform 1 to 7 were taken from reference sequences listed in Table 1. For *MYD88* isoform 8, canonical isoform 2 sequence was taken as template and the corresponding intron sequence between exon 3 and 4 was added. *MYD88* isoform 1 was purchased from Harvard Plasmids (HsCD00296025) and isoform 2 was described earlier [6]. Other CDS were synthesized by the company Genewiz and verified by DNA sequencing. Illustrations were generated using Geneious 5.5.9 and Adobe Illustrator.

*Cell cultures*

All DLBCL cell lines were described previously [6]. DLBCL cell lines were cultured in RPMI supplemented with 20% heat-inactivated fetal bovine serum (FBS, Gibco), except OCI-LY19 (Minimum Essential Medium Alpha, 10% FBS, life technologies). HEK293T cells were cultured in Dulbecco's modified Eagle medium DMEM (Invitrogen) supplemented with 10% FBS. THP-1 WT and MyD88 deficient cells (a gift from V. Hornung, Gene Center, Munich) were cultured in RPMI with 10% FCS.

*Dual Luciferase Assay*

For dual luciferase assays (DLA) 75,000 HEK293T WT or MyD88-deficient I3A cells were plated on a 24-well format and transiently transfected with plasmids expressing *MYD88* isoforms (1-100 ng), firefly luciferase under the NF-κB promoter (100 ng) and Renilla luciferase under a constitutive promoter (10 ng). The total amount of plasmid was adjusted with the empty vector. 48 hours after transfection cells were lysed in passive lysis buffer (Promega) and lysates were measured for luciferase activity on a FluoStar luminescence plate-reader (BMG Labtech). Analysis settings were chosen as recommended in the Dual-Luciferase Reporter Assay System by Promega using MARS data analysis software version 1.20. Graphs and statistics were done in GraphPad Prism version 8.

*SDS-PAGE and immunoblot*

To check expression of MyD88 endogenous proteins and proteins derived from plasmids immunobloting was performed. Cells were lysed in RIPA buffer (20 mM Tris-HCl pH 7.4, 150 mM NaCl, 1 mM EDTA, 10% glycerol, 0.1% SDS, 1% Triton X-100 and 0.5% deoxycholate) supplemented with PhosSTOP, EDTA-free protease inhibitor cocktail (both from Roche) and 0.1 µM PMSF. Reduced and denatured whole cell lysates (WCL) from transient transfected HEK293T cells were separated on 10% Tris-glycine gels using SDS running buffer (25 mM Tris-base, 250 mM glycine and 0.1% SDS) and WCLs used to test endogenous protein were run on 4%–12% gradient gels using MOPS running buffer (Invitrogen). Separated proteins were transferred onto nitrocelullose membranes (GE Healthcare, 0.45 µm). Then, membranes were blocked 1h at room temperature in 5% milk in Tris-buffered saline solution with 0.1% (vol/vol) Tween-20 (TBS-T) and were probed overnight with primary antibodies (all diluted 1:1000): anti-HA H3663 (Sigma-Aldrich), anti-beta-Tubulin 2A1A9 (abcam), MyD88 4D6 (Thermo Fisher), MyD88 D80F5 and 3699 (CST). Next day HRP-conjugated secondary antibodies, anti-rabbit (Biozol) and anti-mouse (Promega), were applied at 1:8000 dilution for 2 h. Membranes were washed three times for 5 min with TBS-T after each antibody incubation. Detection was done by chemiluminescence (Peqlab) and development using a charge-coupled device camera to capture the luminescent signals. Pictures were analyzed and edited in Phusion (Peqlab) and Adobe Illustrator programs.

*Quantitative PCR*

Total RNA isolation was performed by a Qiacube robot using reagents from the RNeasy Mini Kit from Qiagen including DNA digestion (RNase-Free DNase Set, Qiagen). mRNA transcription to cDNA was done manually using High Capacity RNA-to-cDNA (Thermo Fisher). Quantitative PCR was performed in reactions containing 20 ng cDNA, 0.3 or 1 µM of primers, 1x SYBR Green (FastStart Universal SYBR Green Master Rox, Sigma) and RNA-free water. Primers (Table S1) were designed to discriminate all the tested *MYD88* isoforms and map to the exon junctions fulfilling compatibility requirements with SYBR Green mix, which were evaluated in the publicly available software Primer3 (http://bioinfo.ut.ee/primer3-0.4.0/). Of note primers detecting isoform 2 simultaneously amplify isoform 1 (see Fig. S2C); nevertheless isoform 1 suggested to have very low abundance (data not shown). Each sample was analyzed in triplicates in a real-time cycler (Thermo QuantStudio 7 Flex, Thermo Fisher). The cycling profile applied was: 10 min/95 °C; 40 cycles of 95 °C/15 s and 60 °C/1 min, followed by a continuous melt curve stage from 50°C to 95°C. Data was analyzed with the QuantStudio 6 and 7 Flex software and normalized to GAPDH expression.

*Lymphoma dataset analysis*

B cell lymphoma RNAseq libraries from 190 samples, including Burkitt's Lymphoma (BL, n=21), Follicular Lymphoma (FL, n=83), Diffuse Large B cell Lymphoma (DLBCL, n=72), and FL-DLBCL (n=14) were acquired by the German ICGC MMMLSeq consortium and were uploaded as part of several publications to the European genome-phenom archive at EBI: https://www.ebi.ac.uk/ega/home. Naïve B cells (B cells, n=5) and germinal center B cells (GC B cells, n=5) libraries were used as control data and were made public in the same way. Details for library preparation can be found under related papers [45-48]. To visualize alternative splicing events, RNA sequencing data was mapped onto the human reference genome hg38 (UCSC genome) using Segemehl version 2.0 alpha [49]. Splice reads overlapping with the human *MYD88* gene were counted and visualized in Sashimi Plots [50] using R's ggplot2. Also a compositional data approach used in the DIEGO software [51] was applied to analyze differential splicing patterns of the *MYD88* gene. The support number of every splice junction is considered relative to all splice junctions of the *MYD88* gene, and possible variations are analyzed using Wilcoxon's rank sum test as implemented in *R*. Isoform 2 abundance was calculated as 1- *sum of all other splice sites abundances*, because it has no unique splice site. Intron retention was calculated as mean intron coverage relative to the mean coverage of the two flanking exons [52]. DLBCL sub-cluster classification was performed by the ICGC MMML-Seq (https://icgc.org/node/53049; Hübschmann et al, personal communication (MyD88-like n=24 , BCL2-like n=9, BCL6-like n=16 and TP53-like n=19). MyD88 mutations (n=6) M232T, V217F, S219C, I220T, S222R, S243N and T249P were considered as gain-of-function according to Refs. [6,7] apart from L265P (n=5). Generally, tumor cell purity was not adjusted,but there was no significant correlation between tumor cell content and isoform usage. Boxplot graphics were generated using GraphPad Prism version 8.

*CLL dataset analysis*

Chronic Lymphocytic Leukemia (CLL) RNAseq data from 289 patients was acquired from the ICGC-CLL Consortium (https://dcc.icgc.org/releases) and the acquisition and preparation of these libraries has been previously described [12,26]. Quality of raw CLL RNA-seq data in FASTQ files was assessed using ngs-bits:ReadQC (ngs-bits version 0.1 at github.com/imgag/ngs-bits) to identify sequencing cycles with low average quality and base distribution bias. Reads were preprocessed with ngs-bits:SeqPurge [53], mapped using STAR [54] (version 2.5.3a) to the human reference genome GRCh37 (Ensembl) and alignment quality was assessed using ngs-bits:MappingQC. Sashimi plots from CLL data were generated using the Broad Integrative Genome Viewer (IGV, version 2.3.1) to visualize splicing. To reduce false positive hits, only junctions which are covered by at least 5 reads in at least 5 of 289 analyzed samples have been retained. Splice junctions were quantified by normalizing splice junction reads with the total number of spliced reads in the *MYD88* gene. Normalized junction counts were then attributed to one or more matching isoforms.

*Ovarian cancer*

Ovarian cancer RNAseq libraries from 85 patient samples were acquired from the ICGC/OV-AU project (Australian Ovarian Cancer Study, https://dcc.icgc.org/projects/OV-AU). Patient cohort description and libraries preparation can be consulted in previous publications: [28,29]. Ovarian cancer RNA sequencing data was mapped onto the human reference genome hg38

(UCSC genome) and sashimi plots were created using MISO framework version 0.5.3 (https://miso.readthedocs.io) [55].

*Statistical analysis*

Experimental data was analyzed using Excel 2010 (Microsoft) and/or GraphPad Prism 6, 7 or 8 or in *R*, flow cytometry data with FlowJo 10. Normal distribution in each group was always tested using the Shapiro-Wilk test first for the subsequent choice of a parametric (ANOVA, Student's t-test) or non-parametric (e.g. Friedman, Mann-Whitney U or Wilcoxon) test. p-values ($\alpha$=0.05) were then calculated and multiple testing was corrected for in Prism, as indicated in the figure legends. Values <0.05 were generally considered as statistically significant and denoted by * or # throughout. Comparisons were made to unstimulated control, unless indicated otherwise, denoted by brackets.

### Supplemental figure legends

**Supplemental Figure S1.** (A) Mature RNA and amino acid sequences of MyD88 isoforms 2, 4 and 5. Isoform 4 and 5 show out-of-frame translation compared to the canonical TIR domain sequence due to exon 3 skipping. (B) Mature RNA and amino acid sequences of MyD88 isoforms 1 and 2 showing extra amino acids generated by an alternative donor splice site. (C, D) HEK293T cells were transfected with plasmids for signaling incompetent *MYD88* isoforms 3 (C, n=4) and 5 (D, n=4), followed by stimulation of endogenously expressed TLR5 via flagellin and NF-κB dual luciferase assays performed. In C and D one representative of 'n' technical replicates is shown. * = p<0.05 according to two-way ANOVA compared to empty vector (EV).

**Supplemental Figure S2.** (A) Primer design to detect isoforms 1-5. (B) Verification of isoform-specific amplification using plasmid constructs. (C) Immunoblot from lymphoma cell lines and primary B cells with short and long exposure (n=2, red GCB, black ABC). (D) Sashimi plot from GCB cells (n=5). (E) Gating strategy to evaluate B cell purity upon isolation (n=2). (F) Proliferation of stimulated B cells monitored by CFSE (n=2). (G) RT-qPCR analysis of isoforms 3-5 in PBMCs stimulated as indicated (n=3). (H) RNAseq analysis of DLBCL-clusters: MyD88-like (n=24), BCL2-like (n=9), BCL6-like (n=16) and TP53-like (n=19), see Methods. Isoform 2 calculated as 1-(sum of all others). In C, E and F one representative of 'n' biological or technical replicates is shown. G (mean +/-SD) and H (Tukey box and whiskers) represent combined data from n biological or technical replicates. * = $p<0.05$ according to two-way ANOVA (G) and unpaired Student's t-tests (H).

**Supplemental Figure S3.** (A) Mature RNA and amino acid sequences of MyD88 isoforms 2, 6 and 7. Isoform 6 and 7 show a truncated TIR domain. (B) Mature RNA and amino acid sequences of MyD88 isoforms 2 and 8. Isoform 8 shows an early stop-codon compared to the canonical TIR domain sequence due to intron retention. (C) Representative Sashimi plot from CLL samples (n=289). (D) Four representative Sashimi plots from ovarian cancer samples (n = 85). Red shaded boxes point intron retention and orange arcs represents the alternative donor splice site in isoforms 6 and 7.

**Supplemental Table S1: Primers to detect *MYD88* splice isoforms**

| Detection | Forward (5' to 3') | Reverse (5' to 3') | Used concentration | Amplicon size |
|---|---|---|---|---|
| Isoform 1/2 | cccagcattgaggaggattgc | ctcaggcatatgccccaggg | 300 nM | 159 bp |
| Isoform 3 | tgggacccagcattgggc | tccttgctctgcaggtaatc | 300 nM | 247 bp |
| Isoform 4 | atgaccccctgggtgcc | gcacctggagagaggctg | 300 nM | 104 bp |
| Isoform 5 | ggacccagcattggtgcc | gcacctggagagaggctg | 300 nM | 109 pb |
| GAPDH | agccacatcgctcagacac | gcccaatacgaccaaatcc | 1000 nM | 66 bp |

**Figure S1**

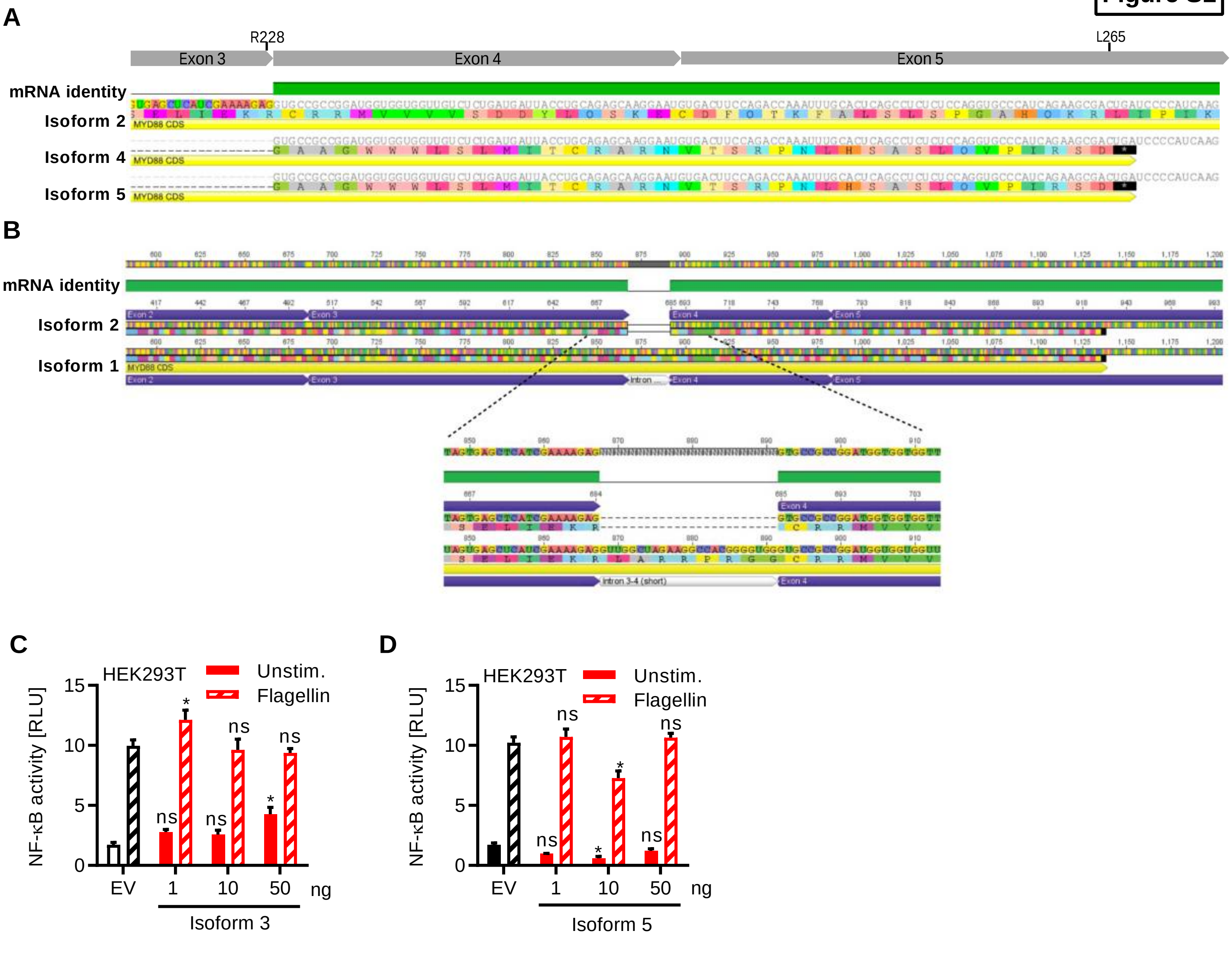
A
R228
L265
Exon 3
Exon 4
Exon 5
mRNA identity
Isoform 2
Isoform 4
Isoform 5
MYD88 CDS
B
mRNA identity
Isoform 2
Isoform 1
Exon 2
Exon 3
Exon 4
Exon 5
Intron
MYD88 CDS
Intron 3-4 (short)
C
HEK293T
Unstim.
Flagellin
NF-κB activity [RLU]
EV
1
10
50
ng
Isoform 3
D
HEK293T
Unstim.
Flagellin
NF-κB activity [RLU]
EV
1
10
50
ng
Isoform 5

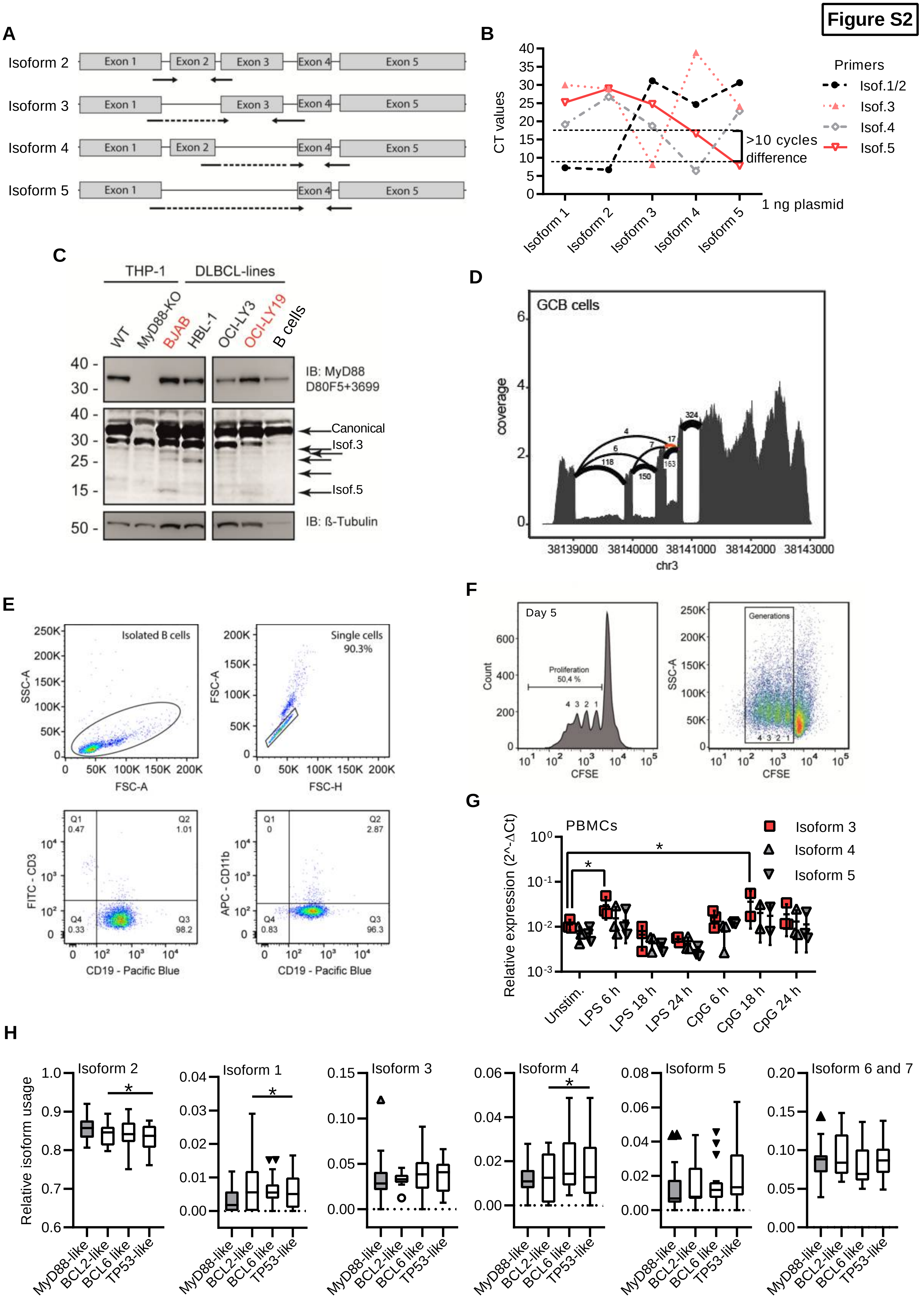

Figure S2
A
Isoform 2
Isoform 3
Isoform 4
Isoform 5
Exon 1
Exon 2
Exon 3
Exon 4
Exon 5
B
CT values
Primers
Isof.1/2
Isof.3
Isof.4
Isof.5
>10 cycles difference
1 ng plasmid
Isoform 1
Isoform 2
Isoform 3
Isoform 4
Isoform 5
C
THP-1
DLBCL-lines
WT
MyD88-KO
BJAB
HBL-1
OCI-LY3
OCI-LY19
B cells
IB: MyD88 D80F5+3699
Canonical
Isof.3
Isof.5
IB: ß-Tubulin
D
GCB cells
coverage
chr3
E
Isolated B cells
Single cells 90.3%
SSC-A
FSC-A
FSC-H
FITC - CD3
APC - CD11b
CD19 - Pacific Blue
F
Day 5
Proliferation 50,4 %
Count
CFSE
Generations
SSC-A
G
PBMCs
Isoform 3
Isoform 4
Isoform 5
Relative expression (2^-ΔCt)
Unstim.
LPS 6 h
LPS 18 h
LPS 24 h
CpG 6 h
CpG 18 h
CpG 24 h
H
Isoform 2
Isoform 1
Isoform 3
Isoform 4
Isoform 5
Isoform 6 and 7
Relative isoform usage
MyD88-like
BCL2-like
BCL6 like
TP53-like

**Figure S3**

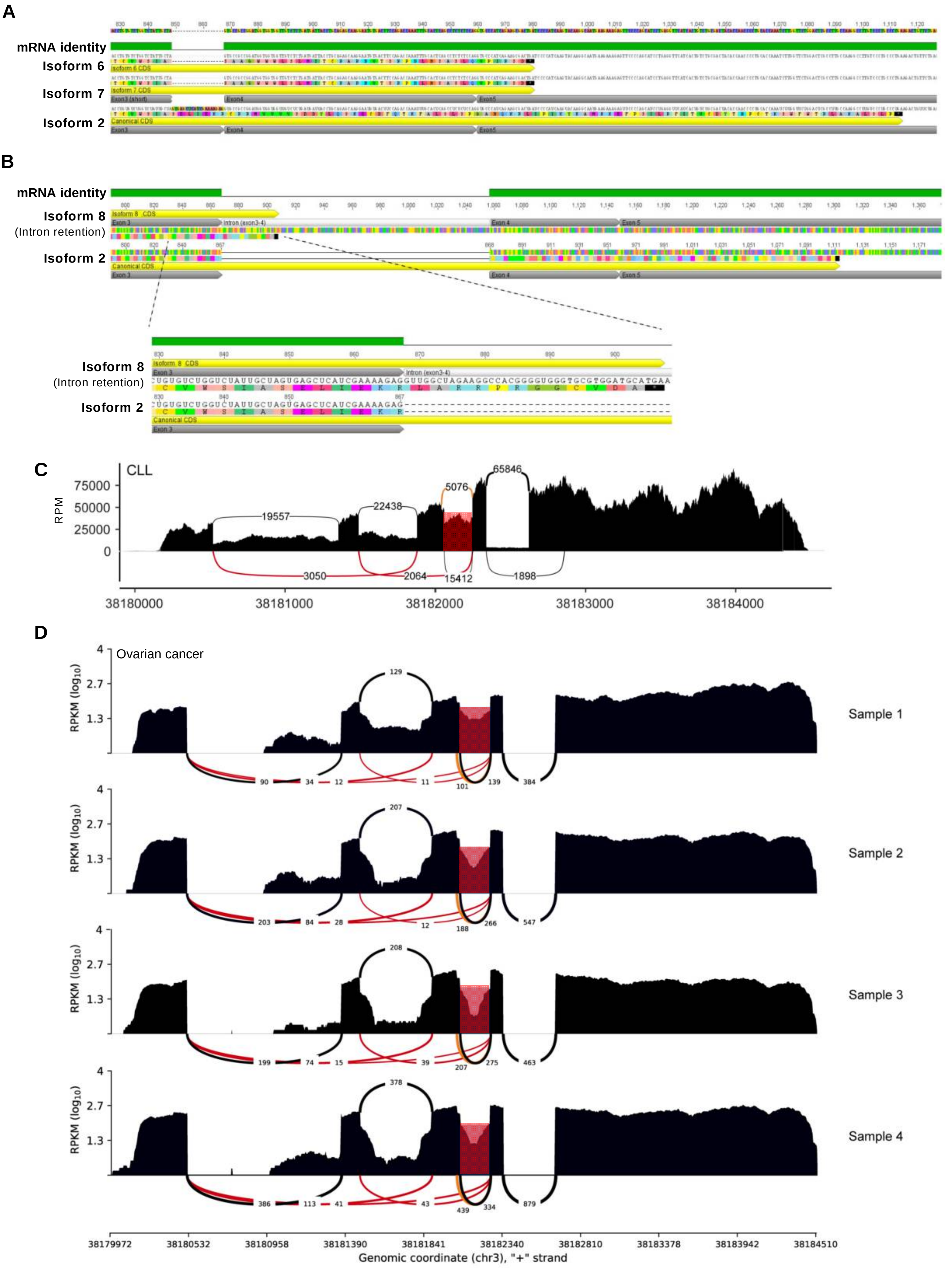
A
mRNA identity
Isoform 6
Isoform 7
Isoform 2
B
mRNA identity
Isoform 8
(Intron retention)
Isoform 2
Isoform 8
(Intron retention)
Isoform 2
CUGUGUCUGGUCUAUUGCUAGUGAGCUCAUCGAAAAGAGGUUGGCUAGAAGGCCACGGGGUGGGTGCGTGGATGCATGAA
CUGUGUCUGGUCUAUUGCUAGUGAGCUCAUCGAAAAGAG
C
CLL
RPM
75000
50000
25000
0
19557
22438
5076
65846
3050
2064
15412
1898
38180000
38181000
38182000
38183000
38184000
D
Ovarian cancer
RPKM (log10)
4
2.7
1.3
129
90
34
12
11
101
139
384
Sample 1
207
203
84
28
12
188
266
547
Sample 2
208
199
74
15
39
207
275
463
Sample 3
378
386
113
41
43
439
334
879
Sample 4
38179972
38180532
38180958
38181390
38181841
38182340
38182810
38183378
38183942
38184510
Genomic coordinate (chr3), "+" strand
